# Trapping studies reveal phenology and reproductive behaviour in two solitary sweat bees (Hymenoptera: Halictidae)

**DOI:** 10.1101/2023.08.09.552673

**Authors:** Alex Proulx, Miriam H. Richards

## Abstract

A transition from univoltine to bivoltine life histories, potentially due to the lengthening of flight seasons, is thought to have preceded the evolutionary transition from solitary to eusocial behaviour in halictid bees. Most behavioral studies which allow us to map these demographic and social transitions on phylogenies have focused on eusocial species, leaving a gap in behavioral data regarding solitary halictid bees. This lack of behavioral trait data is due to the great difficulty in locating the nests of solitary bees for study. However, long-term trapping studies are an effective method providing insights into the demography and social status of halictid bees. We used pan-trapped specimens to investigate flight phenology and solitary reproduction *Lasioglossum leucozonium* (Schrank) and *L. zonulum* (Smith), from the socially and demographically labile subgenus *Leuchalictus*. We found *L. leucozonium* to be univoltine and solitary across its range, regardless of the length of flight season. In contrast, *L. zonulum* is phenologically flexible, being univoltine in the shorter flight seasons of Alberta, but bivoltine in southern Ontario. In both *L. leucozonium* and univoltine *L. zonulum*, higher than expected numbers of old, worn females with declining ovarian development foraged late into the season, suggesting that an indeterminate breeding strategy represents an ancestral condition not found in eusocial species. In bivoltine *L. zonulum*, the large size and higher fecundity of Brood 1 offspring suggest that after evolutionary reversion from eusocial to solitary behaviour, selection for worker-sized daughters may be reversed to selection for larger body sizes. The demographic plasticity demonstrated by *L. zonulum* may hint that partial bivoltinism is more widespread than currently documented. Future studies aimed at clarifying the demographic and social traits of more halictids will pave the way for comparative studies and eventual confidence in tracing how behavioral traits have changed and evolved in halictid bees.

## Introduction

In halictid bees, evolutionary transitions from univoltine to bivoltine or multivoltine life histories seem to be favoured when changing environmental conditions lead to the lengthening of flight seasons, for instance during periods of global warming or during range expansions into warmer climates (Brady et al. 2006). With longer flight seasons, nest foundresses may be able to extend brood production over a longer period than would be possible with shorter flight seasons. Likewise, with longer flight seasons, daughters produced in the first brood have the potential to begin reproduction shortly after eclosion, rather than entering diapause and postponing reproduction until the following year. A switch from univoltine to bivoltine life histories must have preceded evolutionary transitions from solitary breeding to eusociality (Figure 1), so behavioural comparisons of univoltine and bivoltine solitary bees should help to illuminate how this preadaptation for eusociality might have occurred.

**Figure 1:**
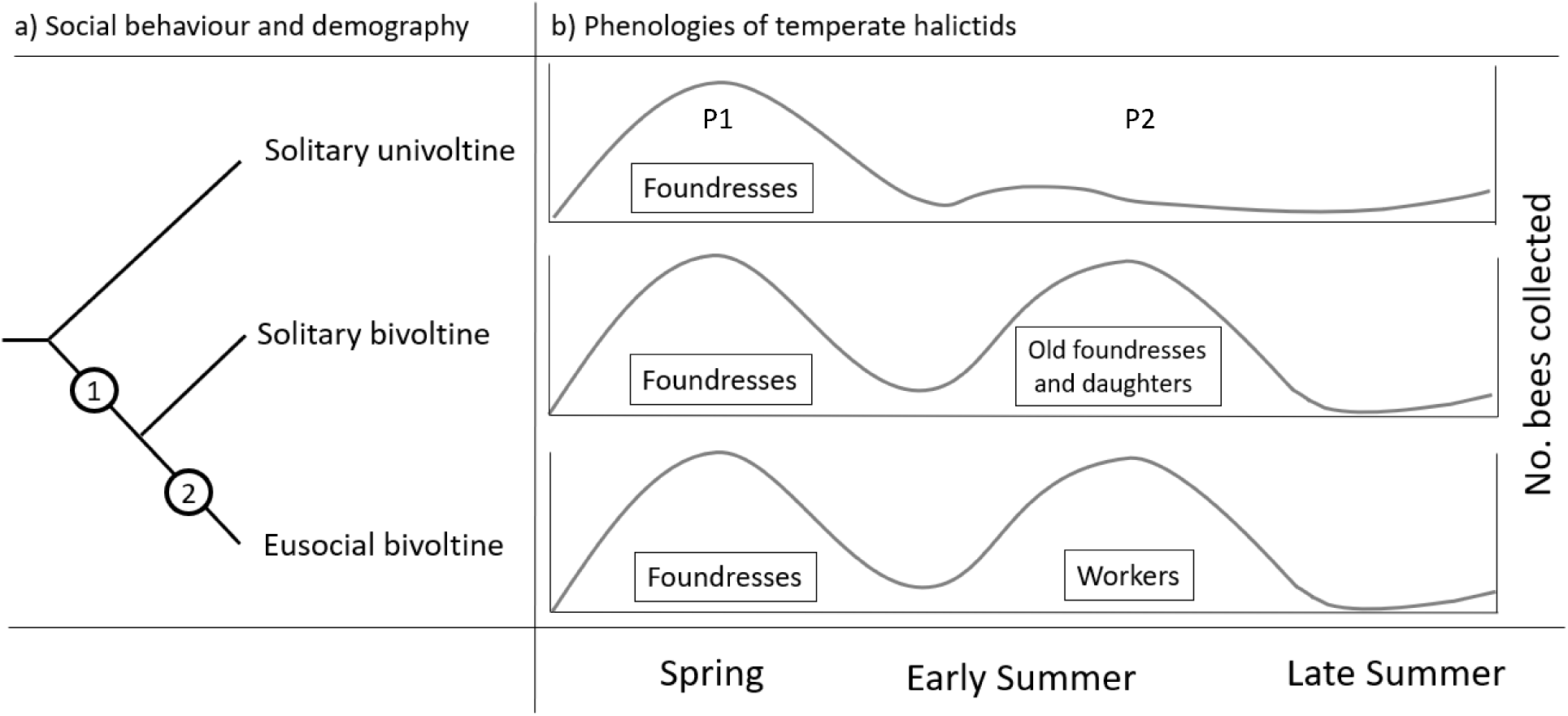
The likely sequence of evolutionary transitions from solitary to eusocial behaviour in sweat bees. **a)** The ancestral solitary univoltine state first shifted demographically from univoltine to bivoltine **(1)** and then shifted behaviourally from solitary to eusocial behaviour **(2)**. **b)** In a trapping study, a univoltine phenology produces a single major peak in collections of foraging females, whereas a bivoltine phenology produces two distinct phases. During Phase 1 (P1), foundresses or queens forage to provision Brood 1. Distinct differences among life histories emerge in Phase 2 (P2).

To reconstruct the steps involved in evolutionary transitions between solitary and eusocial behaviour requires mapping of relevant traits onto detailed phylogenies. But while phylogenetic projects are continuously improving our understanding of sweat bee relationships, behavioural studies have traditionally focussed on eusocial species, and few studies have focussed on solitary species. As a result, behavioural trait data required for reconstruction of transitions between solitary and social states, are sorely lacking. One reason for these gaps in knowledge is that sweat bee reproductive behaviour is best studied by observing female bees at their nests. Since nests can be difficult to find, this creates an impediment to behavioural studies. One solution is to use specimens caught in systematic trapping studies, which can provide considerable insight into a population or species’ mode of reproduction. Trapping studies are especially useful for investigating whether bee populations are univoltine or bivoltine (or even multivoltine). Trapping studies also provide female specimens that can be assessed for traits related to reproduction and social interactions, allowing species that would otherwise remain behaviourally unknown, to be socially categorized as solitary or eusocial.

The large sweat bee genus *Lasioglossum* Curtis is thought to be ancestrally eusocial (Gibbs et al. 2012), which suggests that the reproductive behaviour of extant solitary species could shed considerable light on the steps involved in evolutionary reversals. The subgenus *Leuchalictus* is a particularly interesting clade for studying the relationships between demography and sociality, because it includes a mix of univoltine and bivoltine, as well as solitary and eusocial species (Table 1). The primary objective of this study was to investigate details of phenology and solitary reproduction of two *Lasioglossum (Leuchalictus)* species, *L. leucozonium* (Schrank) and *L. zonulum* (Smith), using specimens collected in pan trapping studies. Both *L. zonulum* and *L. leucozonium* are Holarctic in distribution, and abundant in North America (McGinley 1986; Pesenko et al. 2000).

**Table 1.** Phenology and reproductive behaviour summarized for behaviourally known *Lasioglossum (Leuchalictus).* Type of study refers to observations focussed on females at their nests or females collected on flowers or while foraging. Bivoltinism of *L. majus* in Poland is suggested by the flight phenology presented in Figure 324 of Pesenko et al. (2000).

| Species | Location | Type of study | Phenology | Behaviour | Reference |
| --- | --- | --- | --- | --- | --- |
| *aegyptiellum* (Strand) | Camargue, France | Nests | Bivoltine | Eusocial | Packer 1997 |
| *kansuense* (Bluthgen)<br>(as *L. esoense*) | Hokkaido, Japan | Nests | Univoltine | Solitary | Sakagami et al. 1966 |
| *leucozonium* (Schränk) | Nova Scotia, Canada | Nests | Univoltine | Solitary | Atwood 1933 |
| *leucozonium* (Schränk) | Germany | ? | Univoltine | Solitary | Stockert 1933 |
| *majus* (Nylander) | Lombardy, Italy | Nests | Univoltine | Solitary,<br>communal | Boesi et al., 2009 |
| *majus* (Nylander) | Poland | Flowers? | Bivoltine | Solitary | Pesenko et al. 2000 |
| *mutilum* (Vachal) | Matsue, Japan | Greenhouse nests | Bivoltine | Solitary, eusocial | Miyanaga et al. 1998 |
| *mutilum* (Vachal) | Hachijō-jima Island,<br>Japan | Flowers | Univoltine | Solitary | Takahashi and Sakagami 1993 |
| *occidens* (Smith) | Hachijō-jima Island,<br>Japan | Flowers | Univoltine | Solitary | Takahashi and Sakagami, 1993 |
| *scitulum* (Smith) | Matsue, Japan | Nests | Bivoltine | Solitary, eusocial | Miyanaga et al. 2000 |
| *zonulum* (Smith) | Kyiv, Ukraine | Nests | Univoltine | Solitary | Golubnichaya and Moskalenko<br>1992 |
| *zonulum* (Smith) | Germany | ? | Univoltine | Solitary | Stockert, 1933 |

## Methods

### Collection methods

Specimens were collected during two separate studies of bee communities, one in the Niagara region of southern Ontario (Richards et al. 2011) and the other near Calgary in southern Alberta (Galpern et al. 2017; Vickruck et al. 2019).

In Niagara, bees were collected from 2003-2018 (excluding 2007 and 2014) at multiple sites in St. Catharines, Port Colborne and Wainfleet (42.881° N, 79.378°W to 43.125° N, 79.233°W). Detailed field methods have been previously described (Onuferko et al. 2018; Audet et al. 2021). Briefly, bees were collected in pan traps (170g Solo brand, PS6-0099 plastic bowls, Lake Forest, IL, USA) filled with water and a small amount of detergent (5 drops of Blue Dawn dish soap per litre of water). Thirty traps were laid out along transects (usually linear transects) placed at 3 m intervals, alternating colours (blue, yellow, and white) according to standard protocols (Richards et al. 2011; Onuferko et al. 2018). Traps were laid out by 0900 h and the contents collected after 1500 h (after 1600 h in later years) on days without rainfall. Collections occurred weekly or biweekly, from mid-April or early May until September. Collection weeks were numbered starting with the 16th week of each year (in mid-April). After collection, all bees were stored in 70% ethanol, then pinned and identified using the key from McGinley (1986).

*Lasioglossum zonulum* specimens collected as part of a separate project in southern Alberta were kindly provided by Dr. Paul Galpern, University of Calgary. Collection methods are described in detail in Galpern et al. (2017) and Vickruck et al. (2019). Collections were carried out near Calgary (50.663°N, 113.483°W) between week 9 in mid-June to week 19 in late August in 2016. The agriculturally intensive landscape in this area comprises crop fields and wetlands, with bee sampling beginning after crop seeding had been completed, with collections ending prior to crop swathing. Both pan traps (12 oz. New Horizons, Waldorf, MD) and blue vane traps (SpringStar LLC, Woodinville, WA, USA) were used. These traps were filled with propylene glycol, and samples were collected from the traps at intervals between 5 and 23 days (the average interval was 7.7 ± 2.5 days). In this study, we only included bees collected from pans left out for intervals of two weeks or less, and so the term “week” represents the median date of a collection interval. Specimens were stored in 70% ethanol until pinned and identified by Dr. Lincoln Best using keys listed in the supplementary tables of Vickruck et al. (2019). Vouchers were deposited in the Invertebrate Section of the Museum of Zoology, Department of Biological Sciences, University of Calgary. All sample sizes are given in Table 2.

**Table 2:** Collection information for all three populations of sweat bees examined in this study. The first collections of bees in spring mark the beginning of Phase 1 for each population’s phenology, while the first collections of Brood 1 males indicates the beginning of Phase 2. In Calgary, collections did not begin until Week 9, so bees flying earlier than that would not have been recorded. Niagara collections were pooled to achieve larger samples sizes, and that there were no collections in Niagara in 2007 or 2014.

|  | *L. leucozonium* | *L. zonulum* | *L. zonulum* |
| --- | --- | --- | --- |
|  | Niagara | Calgary | Niagara |
| Collection years | 2003-2018 | 2016 | 2003-2018 |
| No. specimens collected | 197 females, 32 males | 834 females, 13 males | 276 females, 15 males |
| Phenology | Univoltine | Univoltine | Bivoltine |
| Length of flight season | 23 weeks | At least 11 weeks | 20 weeks |
| First bee collections | Week 3 (mid May) | Week 9 <sup>1</sup> | Week 2 (early May) |
| First Brood 1 males | Week 13 (mid-late July) | Week 15 (early August) | Week 12 (mid July) |
| First Brood 1 females | Week 13 (mid-late July) | Week 19 (early September) | Week 13 (mid-late July) |
| Last bee collections | Week 25 (early October) | Week 19 (early September) | Week 21 (mid September) |

<sup>1</sup> In Calgary, collections began in week 9, so earlier bees would have been missed

### Phenology

Strictly speaking, the terms ‘univoltine’, ‘bivoltine’, and ‘multivoltine’ describe life histories that produces one, two, or more generations of offspring per year. However, these terms are also used more loosely to describe sweat bees with annual life histories in which one, two, or more broods of offspring are produced each year, even when these are not necessarily successive generations. These looser definitions are useful (when clarified) because they allow us to use similar terminology for describing the life histories of solitary and eusocial sweat bees.

To evaluate whether bee populations were univoltine or bivoltine, we plotted the average numbers of female bees collected per site per week throughout the flight season. To correct for differences in sampling effort across weeks, weekly bee abundance was estimated as the total number of specimens collected per site per week, pooled across collection years (Onuferko et al. 2018). Specimens collected in Niagara were pooled across years, due to small annual sample sizes. We used non-parametric regressions and a Loess smoother to evaluate when peaks in bee abundance occurred (*ggplot2::geom_smooth* command in R version 4.2.2). Since halictine males do not overwinter, their earliest appearance in collections was used to indicate when spring brood had eclosed as adults, dividing the flight season into a spring phase, P1 (prior to emergence of Brood 1 males) and a summer phase (P2, after male emergence).

### Female reproductive traits and colony social organisation

We measured female traits related to reproductive behaviour and colony social organisation (foundress longevity, body size, wear, and ovarian development), following established methods used in for pinned specimens (Packer et al. 2007; Richards et al. 2010; Corbin et al. 2021). Pinned specimens were examined under 6 to 66x magnification using a stereomicroscope (Zeiss Stemi SV11).

Female size was measured using an ocular micrometer to determine head width (distance across the widest part of the head, including eyes) as well as costal vein length of the forewing. Degree of mandibular wear (MW) is expected to reflect the extent of activities such as digging brood cells; MW was assessed on a scale from 0 (the mandible and apical tooth pointed and in pristine condition) to 5 (mandible completely blunt with no apical tooth). Wing wear (WW) is expected to reflect the extent of flight activity, as bees accumulate wing damage over time (Cartar 1992; Foster and Cartar 2011) and was assessed on a scale from 0 (pristine wing margin along distal edge) to 5 (distal edge destroyed). A total wear (TW) score was calculated by summing MW and WW for a combined score from zero to 10. Females with TW ≤ 1 were considered to be unworn, whereas females with TW ≥ 2 were considered to be worn.

After measuring, pinned specimens were rehydrated in deionized water for ∼24 h and then dissected to assess ovarian development (OD) (Richards et al. 2010; Corbin et al. 2021). Each visible, developing oocyte was assigned a proportional score of 0.25, 0.5, 0.75, or 1, with 1 representing a fully developed oocyte (Richards et al. 2010). These scores were then summed, producing an OD score for each female. Females lacking developed ovaries were assigned OD scores of 0 and those possessing thickened ovaries with no visible oocytes were assigned scores of 0.1. Any female which possessed an oocyte of at least 1/2 size (0.5) was considered “fecund” (Breed 1976). Phase 2 females with TW and OD scores of 0 were assumed to be newly eclosed gynes that had left their nests to mate and feed prior to diapause. All other females were assumed to be foragers collecting provisions for brood.

For all three populations examined, we compared traits of Phase 1 and 2 females, excluding newly eclosed Phase 2 females (Richards et al. 2010; Corbin et al. 2021). Females collected before male emergence were categorized as Phase 1 and those collected afterwards were assigned to Phase 2. Foundress longevity was estimated based on the relative proportions of females collected in Phases 1 and 2, excluding newly eclosed females.

### Statistical analyses

All statistical analyses were conducted using R version 3.6.2 running under R-Studio version 1.2.5033. The package *tidyverse* was used for data curation, manipulation and plotting. The *lm* command was used to produce general linear models. Small annual samples for bees collected in Niagara precluded including annual variation (year effects) in comparisons, despite the potential for annual variation in body size.

## Results

### Lasioglossum leucozonium in Niagara

*L. leucozonium* exhibited a single, extended phase of foraging activity by females (Figure 2). Males and newly eclosed females were first collected during week 13 in early July, suggesting that Brood 1 had reached the stage of adult eclosion. Consequently, week 13 was considered to mark the beginning of Phase 2 (Figure 2). The majority of females (82%) were collected during Phase 1, but worn females continued to be collected until well into autumn (Table 2).

**Figure 2:**
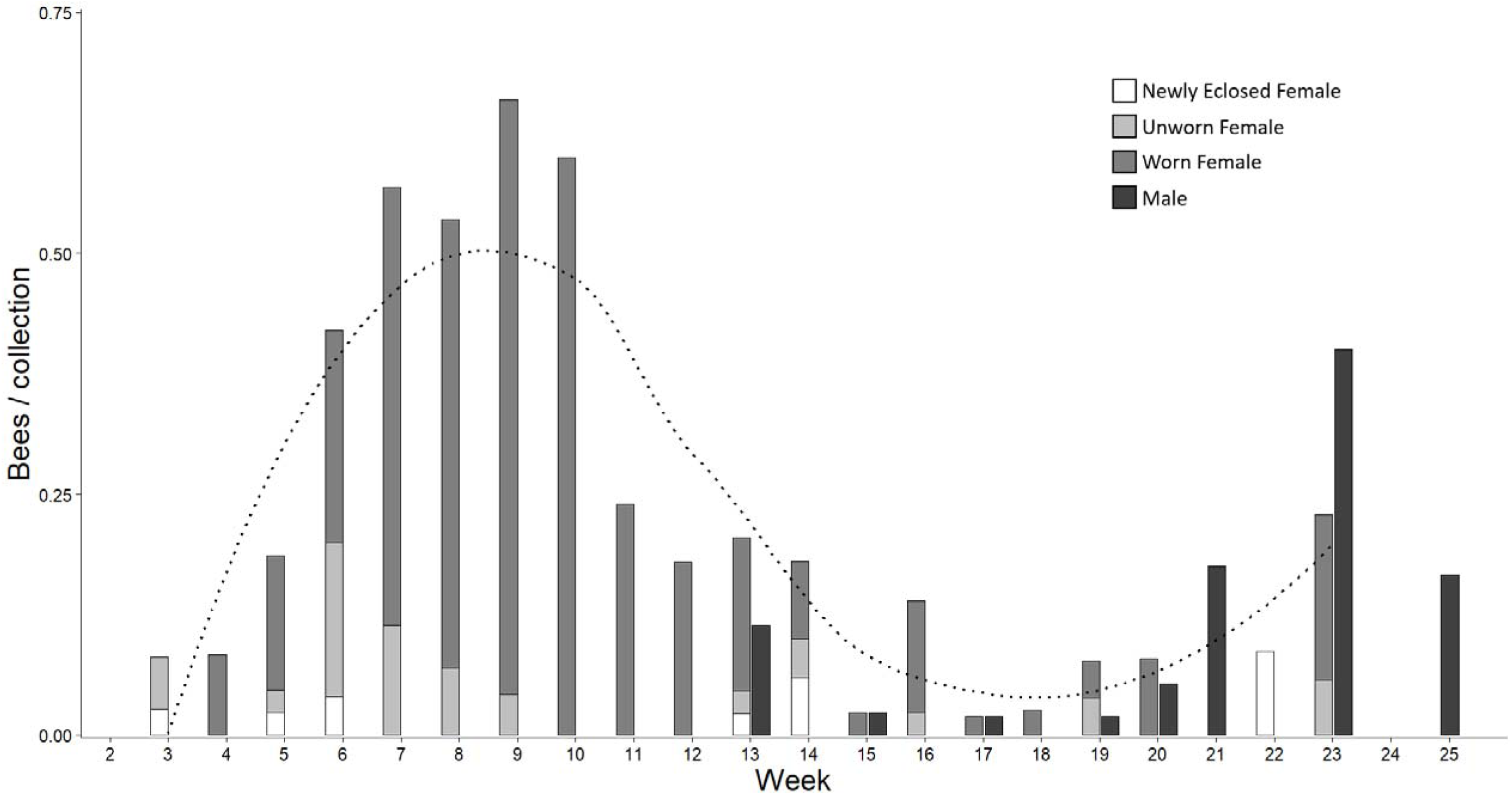
The univoltine phenology of Niagara *L. leucozonium* based on data from 2003 to 2018 pooled. The dotted line represents a Loess smooth fitted to the female data and emphasizes a single major peak in foraging behaviour. Around 78% (n = 153) of female bees were collected in pan traps from weeks 3-12 (early May to early July). Newly eclosed females and males of Brood 1 first appeared in week 13 (early July).

The presence of very worn but fecund females in the P2 samples demonstrates that some foundresses were very long-lived and continued to produce brood throughout the summer.

Traits of *L. leucozonium* females collected in Phases 1 and 2 are compared in Table 3. Phase 1 and 2 females were similar size, as expected if they represented a single cohort. The proportion of worn females was similar during the two phases, but the median wear score was significantly higher, as expected for a single cohort of foundresses that continually accumulated wing and mandibular wear. The proportions of fecund females were also similar, but ovarian scores were significantly lower in Phase 2, suggesting that long-lived foundresses eventually experienced reproductive senescence.

**Table 3:** Traits of adult females from the univoltine population of *L. leucozonium* in Niagara, Ontario. Comparisons of head width and costal vein length are based on Welch two sample t-tests; total wear and ovarian development comparisons are based on Kruskal-Wallis tests (H), and proportions of worn and fecund females are based on Pearson’s X^2^ test with Yates’ continuity correction.

| Female traits | Phase 1 Foundresses | Phase 2 Foragers | Phase 2 Daughters | P1 vs P2 Foundresses |
| --- | --- | --- | --- | --- |
| No. specimens collected | 153 (82.2%) | 33 (17.7%) | 11 |  |
| Head width (mm)<br>(mean ± SD) | 2.42 ± 0.09 | 2.41 ± 0.11 | 2.38 ± 0.10 | t = 0.667, df = 42.183, ns |
| Costal vein length (mm)<br>(mean ± SD) | 3.07 ± 0.11 | 3.03 ± 0.10 | 3.01 ± 0.15 | t = 1.995, df = 49.503, p = 0.051 |
| Total wear score<br>(median, range) | 4 (0 – 9) | 5 (0 – 10) | 0 (0 – 1) | H = 5.46, df = 1, p = 0.019 |
| Proportion worn | 130/149 (87.2%) | 31/33 (93.9%) | | $X^2$ = 0.620, df = 1, ns |
| Ovarian development score<br>(median, range) | 1.25 (0 – 2.75) | 0.75 (0 – 2.5) | 0.1 (0 – 0.1) | H = 4.54, df = 1, p = 0.033 |
| Proportion fecund | 125/153 (81.7%) | 22/33 (66.7%) | | $X^2 = 2.85$, df = 1, ns |

### Lasioglossum zonulum in Calgary

*Lasioglossum zonulum* specimens were collected foraging from week 9 in mid-June to week 19 in late August (Figure 3). Most females (79 %) were collected from mid-to late June (weeks 9-11). The first males were collected on week 16 in early August, which was only 4 weeks before the end of the flight season. As collection week for Calgary was calculated using the median dates of roughly week-long (7.7 ± 2.5 days) collection events, this would suggest that no males were foraging before week 15 at the earliest. A single, newly eclosed female (TW = 0, OD = 0) was collected in late August (week 19). As there was a single major phase of foraging activity weeks 9 to 11 prior to the emergence of Brood 1, *L. zonulum* is univoltine in southern Alberta.

**Figure 3:**
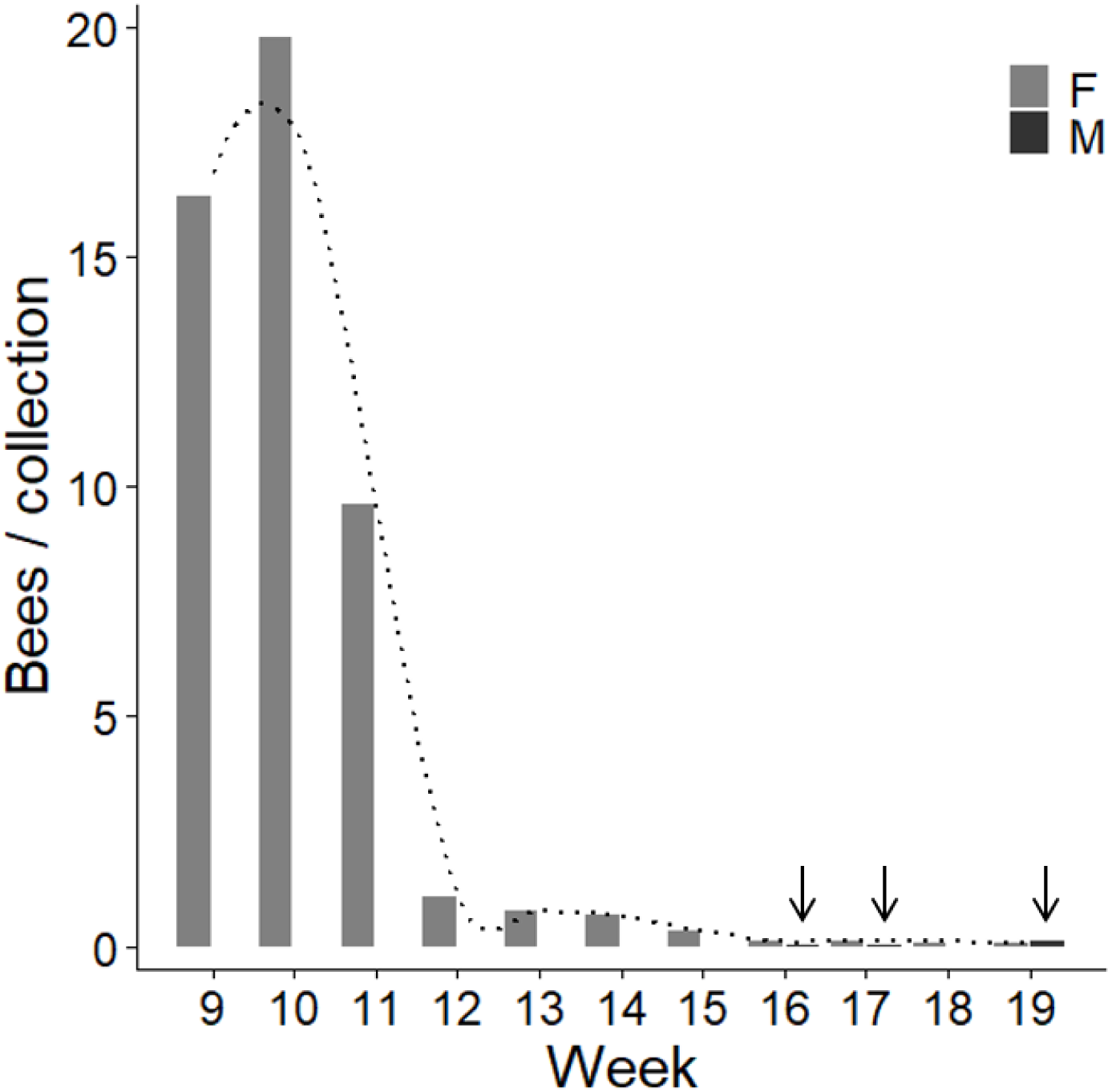
The univoltine phenology of Calgary *L. zonulum* collected in 2016 with a Loess smoother fitted emphasizes a single major peak in female foraging behaviour. Females are shown in light grey, while males are shown in dark grey and emphasized with arrows. Males only made up 1.5% (13/847) of collected bees and were only collected after week 16.

Females collected before and after Brood 1 emergence are compared in Table 4. About 21% of foundresses were collected during Phase 2, suggesting that most females die before the emergence of their oldest brood, but a few long-lived females forage and provision brood right up to the end of the flight season. As predicted for a single cohort of foundresses, P1 females were similar in size. The proportions of unworn females were similar in the two phases, but P2 females had markedly higher wear scores. The proportions of fecund females were similar in both phases, but ovarian scores were significantly lower during Phase 2. As for *L. leucozonium*, these patterns support the hypothesis that a single cohort of foundresses provisions brood each year.

**Table 4:** Traits of adult females from the univoltine of *L. zonulum* in Calgary, Alberta. Comparisons of head width and costal vein length are based on Welch two sample t-tests; total wear and ovarian development comparisons are based on Kruskal-Wallis tests (H), and proportions of worn and fecund females are based on Pearson’s X^2^ test with Yates’ continuity correction.

| Female traits | Phase 1<br>Foundresses | Phase 2<br>Foragers | Phase 2<br>Daughters | P1 vs P2<br>Foundresses |
| --- | --- | --- | --- | --- |
| No. specimens | 108 (79.4%) | 28 (20.6%) | 1 |  |
| Head width (mm)<br>(mean ± SD) | 2.58 ± 0.09 | 2.57 ± 0.09 | 2.41 | t = 0.738, df = 41.350, ns |
| Costal vein length (mm)<br>(mean ± SD) | 3.27 ± 0.11 | 3.26 ± 0.12 | 3.08 | t = 0.373, df = 39.595, ns |
| Total wear score<br>(median, range) | 4 (1–10) | 9 (4–10) | 0 | H = 48.157, df = 1, p = 3.93e-12 |
| Proportion worn | 101/108 (93.5%) | 28/28 (100%) | | $X^2$ = 0.816, df = 1, ns |
| Ovarian development score<br>(median, range) | 0.50 (0–2.25) | 0.1 (0–0.1) | 0 | H = 10.116, df = 1, p = 0.00147 |
| Proportion fecund <sup>2</sup> | 58/98 (59.2%) | 10/28 (32.5%) | | $X^2$ = 0.816, df = 1, ns |

<sup>2</sup> Proportion of dissected females with at least one 1/2-size oocyte

### Lasioglossum zonulum in Niagara

The flight season of Niagara *L. zonulum* lasted 20 weeks from early May to early September (Figure 4). There were two major periods of foraging activity, separated by a slight hiatus in activity during weeks 9-10, with almost exactly equal numbers of female specimens being collected in Phases 1 and 2 (Table 5). Newly eclosed females from Brood 1 were first collected in early July on week 12, while the first males were collected in week 13. As the appearance of newly eclosed females preceded that of males, week 12 was taken as the beginning of Phase 2. The bivoltine phenology of *L. zonulum* in Niagara contrasts strongly with the univoltine phenology in Calgary and demonstrates that this species is behaviourally flexible for this trait.

**Figure 4:**
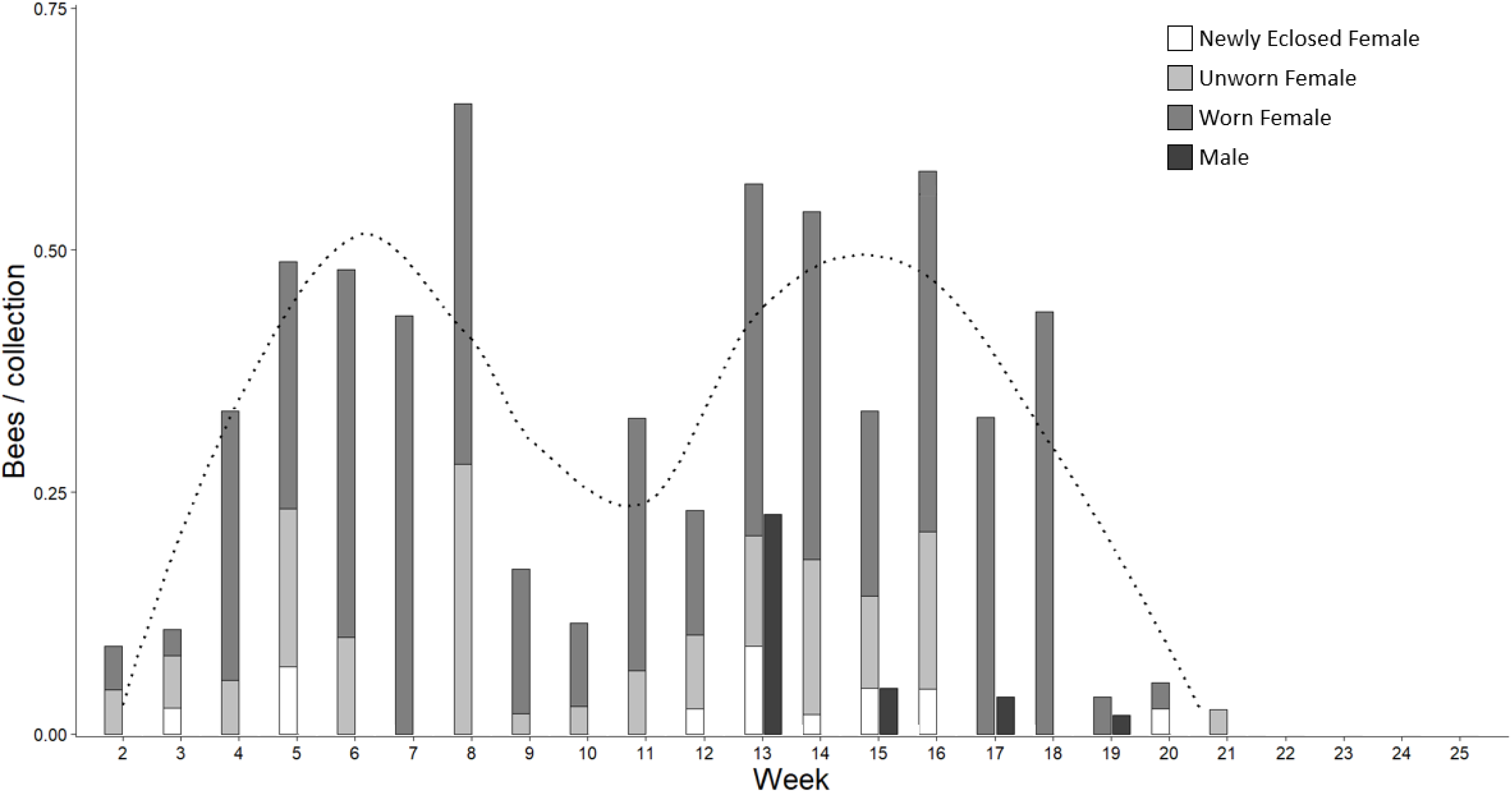
The bivoltine phenology of Niagara *L. zonulum* based on data pooled across 2003-2018. The dotted line represents a Loess smoother fitted to the female data and emphasizes two phases in foraging behaviour. Note the decline in captures of females in late June between weeks 9 and 12. Newly eclosed females from Brood 1 first appeared in week 12, and male bees in week 13 (early July).

**Table 5:** Traits of adult females for the bivoltine population *L. zonulum* in Niagara, Ontario. Comparisons of head width and costal vein length are based on Welch two sample t-tests; total wear and ovarian development comparisons are based on Kruskal-Wallis tests (H), and proportions of worn and fecund females are based on Pearson’s X^2^ test with Yates’ continuity correction.

| Female traits | Phase 1<br>Foundresses | Phase 2<br>Foragers | Phase 2<br>Daughters | P1 vs P2<br>Foundresses |
| --- | --- | --- | --- | --- |
| No. specimens | 137 (55.2%) | 111 (44.8%) | 27 |  |
| Head width (mm)<br>(mean ± SD) | 2.48 ± 0.11 | 2.52 ± 0.10 | 2.47 ± 0.13 | t = -2.50, df = 243.15, p = 0.013 |
| Costal vein length (mm)<br>(mean ± SD) | 3.09 ± 0.12 | 3.12 ± 0.10 | 3.08 ± 0.12 | t = -1.93, df = 267.17, p = 0.054 |
| Total wear score<br>(median, range) | 2 (0–10) | 5 (0–10) | 1 (0–1) | H = 47.36, df = 1, p < 5.91e-12 |
| Proportion worn | 108/137 (78.8%) | 101/111 (91.0%) | 0% | $X^2$ = 5.95, df = 1, p = 0.0147 |
| Ovarian development score<br>(median, range) | 0.75 (0–3) | 1.5 (0–3.25) | 0 (0–0.1) | H = 18.36, df = 1, p = 1.82e-05 |
| Proportion fecund <sup>3</sup> | 68 / 137 (49.6%) | 93 / 111 (83.8%) | 0% | $X^2$ = 29.92, df = 1, p = 4.51e-08 |

Traits of P1 and P2 females are compared in Table 5. Phase 1 females were deemed to be nest foundresses, while Phase 2 females likely represented a mix of old foundresses, Brood 1 females provisioning Brood 2, and newly eclosed daughters, mostly like produced in Brood 2. About 55% of foragers collected were from Phase 1 and about 45% were from Phase 2. P2 females were significantly larger, suggesting that Phase 1 foundresses had provisioned female brood larger than themselves. P2 females were also significantly more worn and had higher ovarian scores. This suggests that in this bivoltine population, per capita brood production may have been higher in Phase 2 (summer) than in Phase 1 (spring).

## Discussion

*Lasioglossum leucozonium* was found to be univoltine and solitary in southern Ontario, a region where the bee flight season is long enough for other *Lasioglossum* to be bivoltine (Awde and Richards 2018; Corbin et al. 2021). Previous studies have found *L. leucozonium* to be univoltine in other portions of its range, as well (Table 1), so this species seems to be obligately univoltine and solitary. In contrast, we found that *L. zonulum* exhibits flexible phenology, being bivoltine in southern Ontario, but univoltine in Alberta and other areas (Table 1). Thus, *L. zonulum* is facultatively bivoltine.

### Traits of univoltine solitary sweat bees

Our examination of female traits in the univoltine populations of *L. leucozonium* and Calgary *L. zonulum* demonstrated some remarkable similarities. In both, about 80% of foundresses were collected during Phase 1, suggesting that in univoltine populations, most females die or at least, cease brood provisioning, prior to the emergence of their brood as adults. However, about 20% of foundresses were collected in Phase 2, so quite a substantial proportion of foundresses were still engaged in provisioning activity after the emergence of the oldest spring brood. In fact, very long-lived foundresses might have produced offspring over a period as long as 5 months (May to September, Table 2). The similar size distributions of Phase 1 and 2 females, together with higher wear and lower ovarian development in Phase 2, are consistent with the idea that most females collected are indeterminate breeders that begin brood production in spring and continue to provision and lay eggs until senescence and morbidity catch up with them. For indeterminate breeders, the major predictor of maternal reproductive success is maternal longevity (Bosch and Vicens 2006).

Indeterminate breeding likely represents the ancestral maternal condition in bees and many other insects that are solitary breeders. In contrast, subsocial and eusocial bees exhibit determinate breeding, with females terminating egg-laying long before becoming senescent. In subsocial and social carpenter bees with extended maternal care, mothers feed newly eclosed offspring in a second bout of offspring provisioning (Hogendoorn and Velthuis 1999; Hogendoorn et al. 2001). In these bees, it appears that mothers produce fewer eggs than they could, ensuring their own survival to provide the extended maternal care that their brood will require after reaching adulthood. In eusocial sweat bees, foundresses (queens) provision a limited number of brood, then cease egg-laying while they wait for their daughters to reach adulthood and become workers (Schwarz et al. 2007). Thus queens limit the size of the worker brood, suggesting determinate production of Brood 1. This suggests that evolutionary transitions from solitary to subsocial or eusocial behaviour must involve a switch from indeterminate and determinate brood production.

Obligately univoltine species like *L. leucozonium* are generally assumed to be obligately solitary, but the long lifespans of some foundresses create opportunities for behavioural interactions among foundresses and their adult daughters. Such interactions might influence whether daughters overwinter in their natal nests or construct hibernacula elsewhere. Overwintering location, in turn, could influence the probability of communal nesting, because foundresses that overwinter in the same nest might be more likely to remain together in spring (Packer and Knerer 1985). Communal behaviour likely occurs in most solitary sweat bees, at least occasionally, including *L. leucozonium* (McGinley 1986), but cannot be distinguished from solitary nesting without extensive observations of females at their nests.

### Traits of bivoltine solitary sweat bees

The Niagara population of *L. zonulum* was bivoltine, so this species is phenologically flexible, bivoltine in the longer summers of southern Ontario but univoltine in the shorter flight seasons of Alberta and other locations (Table 1). Several other members of the subgenus are facultatively bivoltine. *Lasioglossum majus* (Nylander) is mainly solitary in northern Italy (Boesi et al. 2009), but data presented by Pesenko (2000, Figure 324) suggest that it is bivoltine in Poland. *Lasioglossum mutilum* (Vachal) is facultatively eusocial and produced a small number of eusocial colonies in a bivoltine greenhouse population (Miyanaga et al. 1998). Eusociality in *Lasioglossum aegyptiellum* (Strand) is known from observations of a single colony (Packer 1997) and although usually classified as eusocial, could also be facultatively eusocial. We initially anticipated that *L. zonulum* might also provide evidence of eusocial behaviour in Phase 2. In eusocial sweat bees, Phase 2 females (workers) generally are smaller, less worn, and have lower ovarian development than Phase 1 females (queens) (Packer et al. 2007; Richards et al. 2010; Corbin et al. 2021). The opposite patterns were found in Niagara *L. zonulum*; Phase 2 females were larger, more worn, and had higher ovarian development than Phase 1 females, which convincingly rules out facultative eusociality in this species.

The significantly larger sizes and higher ovarian development scores of Phase 2 females in Niagara *L. zonulum* seem to be unique observations for bivoltine sweat bees. In female bees, larger body size is associated with higher fecundity, longer flight distances and more efficient foraging, and enhanced overwintering survival (Bosch and Vicens 2006; Zurbuchen et al. 2010; Rehan and Richards 2010). In other biovoltine sweat bees both solitary and eusocial, foundresses (Phase 1 females) are generally larger than the daughters that become Phase 2 foragers, with a proportional size difference between generations similar to that observed in eusocial species. For instance, in *L. (Hemihalictus) villosulum* (Kirby), foundresses were about 7.4% larger than daughter generation, based on wing length (Plateaux-Quénu et al. 1989), and in *L. (Dialictus) ellisiae* (Sandhouse), Phase 1 females were about 3.5% larger, based on head width (Corbin et al. 2021). Given the advantages of large size for female bees, perhaps it is not the large size of *L. zonulum* daughters that begs explanation, but the small size of daughters in other bivoltine solitary species. Perhaps bivoltine solitary species with small daughters represent relatively recent reversions from eusociality to solitary behaviour, and so the size difference between foundresses and their daughters reflects ancestral size differences between queens and workers. This might imply that species like *L. zonulum* are ancestrally solitary or represent older reversions to solitary behaviour, in which the selective advantages of large size for daughters have produced the current size distribution.

### Two kinds of solitary phenology

A major difference between univoltine and bivoltine physiology lies in diapause. In univoltine sweat bees, such as *L. leucozonium*, newly eclosed females soon enter overwintering diapause prior to founding nests and raising brood. In bivoltine species, Brood 1 females eclose and almost immediately begin their own brood production, omitting diapause altogether, whereas Brood 2 females enter diapause preparatory to becoming foundresses in the following spring. The flexibility to omit or repeat diapause allows sweat bees to adjust their life history in response to local environmental circumstances. Evidently, *L. zonulum* can now be added to the list of species displaying geographic variation in phenology. The most dramatic example is provided by *L. (Sphecodogastra) lusorium* (Cresson) [studied as *Evylaeus galpinsiae* (Bohart and Youssef 1976)], a floral specialist that is completely dependent on primroses (*Oenothera*) for brood pollen provisions. In some years, this plant flowers in abundance and in other years it flowers not at all. Remarkably, when flowers are unavailable, *L. lusorium* females can re-enter diapause for a second winter, becoming foundresses the following spring. Clearly diapause flexibility is key to extension of life histories from univoltine to multivoltine reproduction. It must also be key for the shortening of life histories from multivoltine to univoltine life histories, as illustrated by *L. zonulum* which we found to be bivoltine in southern Ontario, but univoltine in Calgary, as well as *L. majus* and *L. mutilum* (Table 1).

Geographic variation in voltinism suggests that in bivoltine or multivoltine sweat bees, all newly eclosed females have the potential to enter diapause or to begin reproduction shortly after eclosion. Given this flexibility, it seems likely that many, perhaps all, bivoltine species, are actually partially bivoltine (Yanega 1988). In partially bivoltine species, some Brood 1 females may mate and enter diapause before becoming foundresses the following spring instead of raising their own brood during Phase 2 of the nesting cycle. Given the bet-hedging advantages to mothers of producing some Brood 1 daughters that diapause while their sisters initiate reproduction immediately (Seger 1983), partial bivoltinism is likely widespread. Direct evidence of partial bivoltinism can only be obtained from multi-year observational studies of marked females at their nesting sites (Yanega 1989).

## Conclusions

In sweat bees, evolutionary transitions to bivoltinism or multivoltinism represent a necessary preadaptation in the evolution of eusociality, because eusocial colonies are virtually always annual, with spring queens producing a first brood of workers that then help their mother to raise a subsequent ‘reproductive’ brood (Figure 1) (Schwarz et al. 2007). Thus transitions from solitary to eusocial behaviour appear to be constrained to a particular evolutionary sequence involving a transition from univoltine to bivoltine phenology, followed by a transition to eusociality (Figure 1a). An open question is whether evolutionary reversals from eusocial to solitary behaviour recapitulate in reverse the sequence of events in solitary-to-eusocial transitions and whether secondarily solitary species might retain certain behavioural traits from eusocial ancestors that distinguish them from ancestrally solitary species (Lewis and Richards 2017). Comparisons among bivoltine solitary bees might eventually shed some light on this question. A detailed phylogeny of behaviourally known halictid bees is desperately needed so that we can stop speculating about how social traits evolve and begin to confidently trace trait changes that occur in evolutionary transitions between solitary and eusocial behaviour.

## Acknowledgments

We are very grateful to Paul Galpern for providing specimens from Calgary and Lincoln Best for identifying them. We also thank members of the Brock Bee Lab for their help and encouragement, especially while AP finished his thesis during the covid pandemic.

